# Specifying precision in visual-orthographic prediction error representations for a better understanding of efficient reading

**DOI:** 10.1101/2024.02.29.582776

**Authors:** Wanlu Fu, Benjamin Gagl

## Abstract

Efficient visual word recognition presumably relies on orthographic prediction error (oPE) representations. Based on a transparent neurocognitive computational model rooted in the principles of the predictive coding framework, we postulated that readers optimize their percept by removing redundant visual signals, allowing them to focus on the informative aspects of the sensory input (i.e., the oPE). Here, we explore alternative oPE implementations, testing whether increased precision by assuming all-or-nothing signaling and more realistic word lexicons results in adequate representations underlying efficient word recognition. We used behavioral and electrophysiological data (i.e., EEG) for model evaluation. More precise oPE representations (i.e., implementing a binary signaling and a frequency-sorted lexicon with the 500 most common five-letter words) explained variance in behavioral responses and electrophysiological data 300 ms after stimulus onset best. The original less-precise oPE representation still best explains early brain activation. This pattern suggests a dynamic adaption of represented visual-orthographic information, where initial graded prediction errors convert into binary representations, allowing accurate retrieval of word meaning. These results offer a neuro-cognitive plausible account of efficient word recognition, emphasizing visual-orthographic information in the form of prediction error representations central to the transition from perceptual processing to the access of word meaning.

## Introduction

The predictive processing framework of cortical functioning assumes that the brain uses learned patterns in the environment to optimize information processing starting from the sensory input (Srinivasan, Laughlin, & Dubs, 1982; De Lange, Heilbron, & Kok, 2018; Rao & Ballard, 1999). The framework assumes that by making predictions based on past experiences, the brain constructs probabilistic inferences about upcoming sensory information (Clark, 2013). The predominant paradigm for investigating predictive processing is to manipulate predictability from the immediate or recently learned context of sensory input (Hawelka, Schuster, Gagl, & Hutzler, 2015; Heilbron, Richter, Ekman, Hagoort, & de Lange, 2020; Hofmann, Remus, Biemann, Radach, & Kuchinke, 2022; Eisenhauer, Gagl, & Fiebach, 2022; Yan, de Lange, & Richter, 2023; Alink, Schwiedrzik, Kohler, Singer, & Muckli, 2010). Recently, we showed that humans not only implement predictive processing based on the immediate context but also based on predictable patterns that can be derived from our long-term memory (Gagl et al., 2020). Here, we investigate the influence of algorithmic and knowledge-based precision by implementing the orthographic prediction error in multiple variants to allow an investigation of how prediction error representations are formed. We implement alternative hypotheses based on transparent computational models (Guest & Martin, 2021) and evaluate them based on human behavior and brain activation.

The principles of predictive coding (Srinivasan et al., 1982; Rao & Ballard, 1999) provide a framework for investigating the integration of sensory information (i.e., vision) and stored knowledge essential to reading (i.e., phonology, semantics). The Prediction Error Model of Reading (PEMoR, Gagl et al., 2020) postulates that this integration is based on the removal of redundant visual signals to focus on the informative aspects of the percept (i.e., orthographic prediction error; oPE). Thus, we believe that the oPE representation, which integrates sensory and stored information, is essential for efficient word perception.

A central characteristic of the PEMoR is its simplicity, allowing an implementation that is fully transparent, without free parameters based only on a single image computation. Here, we exploit that simplicity to investigate two aspects of precision in predictive processing. In general, predictive processing becomes more precise when the upcoming information can be better predicted, resulting in a reduction of uncertainty (Clark, 2013). High-quality visual input is crucial for the reading process, as it allows the brain to fine-tune its response to characteristics of sensory information, allowing efficient decoding by a reduction of uncertainty and a focus on what information is most important for visual word recognition (Frost & Katz, 1989; Price & Devlin, 2011; Grainger, 2018; Woolnough et al., 2021). High precision allows the brain to quickly and accurately decode discrepancies between expected and actual visual input, leading to more efficient error correction and adaption during reading (Legge, Rubin, Pelli, & Schleske, 1985; Pelli, Burns, Farell, & Moore-Page, 2006; Rauschecker et al., 2011; Jordan, Dixon, McGowan, Kurtev, & Paterson, 2016; Gagl et al., 2020). In addition, high precision, in a predictive context, is associated with more effective visual orthographic processing facilitating word recognition (Tamminen & Gaskell, 2013; Elgort, Brysbaert, Stevens, & Van Assche, 2018; Eisenhauer, Fiebach, & Gagl, 2019; Gagl et al., 2020). Furthermore, well-described predictability and priming effects further suggest that precision plays a crucial role in reading, where it not only decodes the visual form of words but also helps to combine orthographic information with the semantic and context information (Dambacher, Kliegl, Hofmann, & Jacobs, 2006; Dimigen, Sommer, Hohlfeld, Jacobs, & Kliegl, 2011; Gagl, Hawelka, Richlan, Schuster, & Hutzler, 2014; Carreiras, Armstrong, Perea, & Frost, 2014; Brust & Denzler, 2019; Eisenhauer et al., 2022). In other words, precision in early visual-orthographic word processes precedes effective extraction of meaning from text.

Here, we explore two algorithms that implement a binary all-or-nothing prediction error representation that increases precision in visual orthographic representation by a shift from a graded error for each pixel (i.e., any value between 0 and 1) to a binary (error or not) representation. One algorithm implemented the binary prediction error on the level of the error (see Fig. 1A) and one on the level of the prediction (see Fig. 1B). These hypotheses implement a precision increase by a simplification of the representation from graded to binary prediction errors, potentially more likely reflecting signals implemented in neuronal all-or-nothing signal (i.e., spikes; Qin et al., 2020; Saszik & DeVries, 2012). In the original PEMoR, the prediction results from a pixel-by-pixel mean that integrates the visual appearance of all words stored in the mental lexicon. Thus, the resulting prediction and prediction error representations comprised graded values for each pixel. When assuming binary all-or-nothing coding, the value of a pixel can be either 0 or 1 (see Fig. 1).

**Figure 1:**
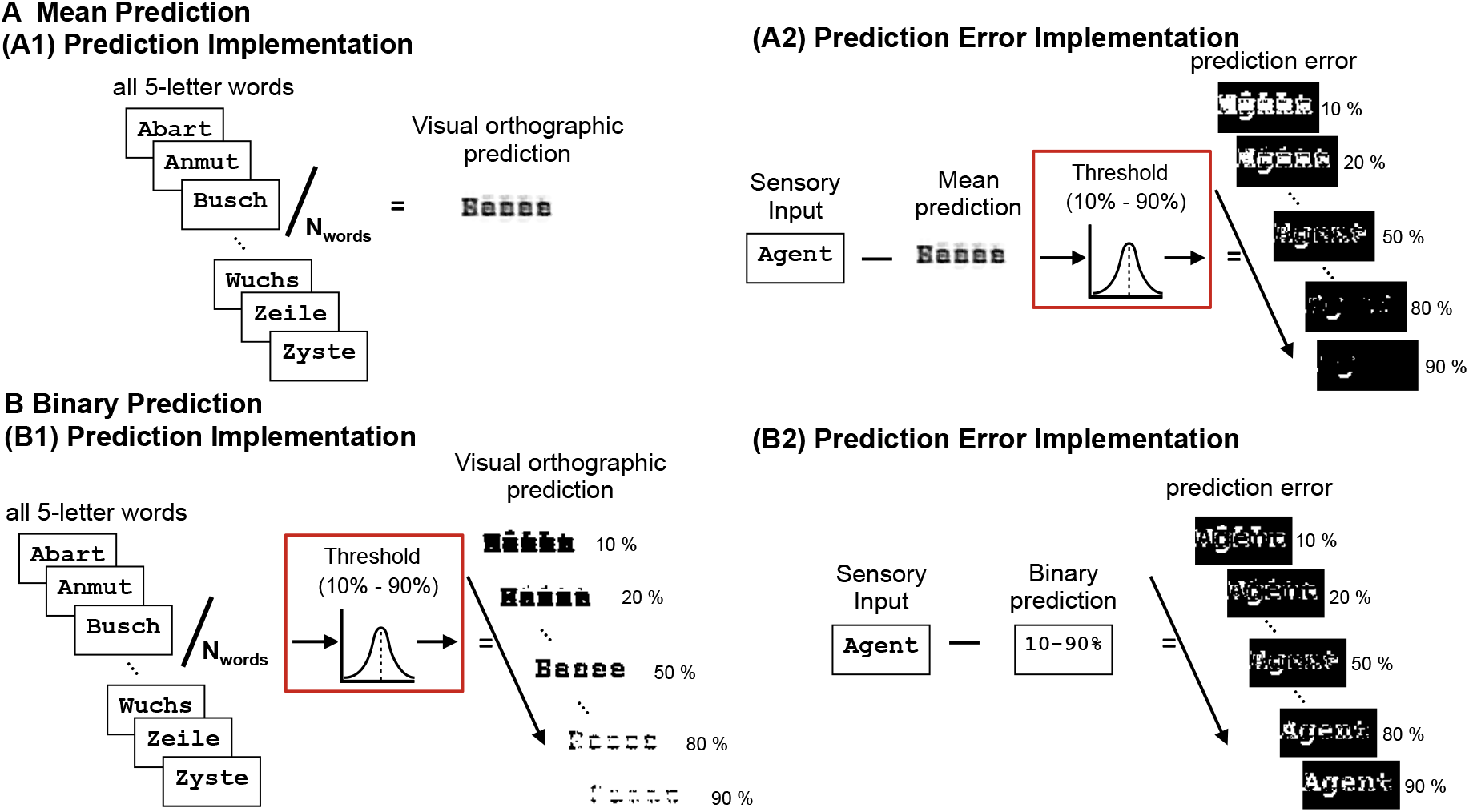
Schematic depiction of the PEMoR adaptations to increase precision in the prediction error representations. (A) PEMoR version that assumes a graded prediction (i.e., based on a pixel-by-pixel mean calculation combining all words stored in the lexicon, i.e., the redundant visual information; see A1) and a binary prediction error output after a threshold implemented from the integration of the prediction and the sensory input (see A2). After the redundant visual information is removed from the sensory input (i.e., by subtraction), we implemented thresholds for calculating stimulus-specific prediction error (i.e., from 10-90% in steps of 10%). (B) The second new PEMoR version assumes a threshold to achieve a binary prediction that automatically results in a binary output. Again, we implemented the knowledge-based prediction but then implemented a threshold to get a binary prediction (i.e., with the same thresholds as above; see B1). As a consequence, after removing the new binary prediction from sensory input (i.e., that is, monochrome), the orthographic prediction error representation is automatically a binary representation but with different representations for the different thresholds (e.g., cp. examples from A2 and B2).

The first hypothesis for the present study implements a mean prediction similar to the original PEMoR implementation (Fig. 1A). In addition, to increase the precision of the prediction error, a threshold is administered after we calculate the prediction error, resulting in a precise binary prediction error representation. In contrast, we achieve higher precision in the second hypothesis by implementing a threshold already at the prediction, resulting in a binary instead of a graded signal (Fig. 1B). Integrating the binary prediction with the binary (i.e., monochrome) sensory input results in a more precise prediction error. Thus, here, we achieve increased prediction error precision based on two different algorithms, implementing a threshold either at the level of the prediction or after calculating a graded prediction error.

Furthermore, we test one additional source that could increase precision in the prediction error representation: the number and type of words included in the lexicon of the model. Numerous studies have shown that contextual factors influence predictive processing in language comprehension (Griffiths, Chater, Kemp, Perfors, & Tenenbaum, 2010; Alink et al., 2010; Kok, Jehee, & De Lange, 2012; De Lange et al., 2018; Heilbron & Chait, 2018; Zhang, Zhao, & Wang, 2020; Shain, Blank, van Schijndel, Schuler, & Fedorenko, 2020; Schrimpf et al., 2021; Eisenhauer et al., 2022). The PEMoR, in its current implementation, has the context of a word recognition task that includes words with a fixed number of letters (i.e., five letters). Thus, in principle, one must perceive the visual stimulus and map it to an entry in the lexicon (i.e., the memory for learned words). Here, by assuming an active predictive word recognition process, lexicon items are the stored knowledge used to predict the upcoming sensory information (see Fig. 1). Therefore, in addition to increasing the precision on the algorithm level (see above), we can also increase the precision by assuming different lexicons, including fewer words than assumed in the original implementation (Gagl et al., 2020).

Restricting the stimulus material to only five-letter words increases the precision by better predicting where the stimulus is presented, removing artificial noise that is the consequence of unavailable parafoveal preview in word recognition tasks. In natural reading, we reliably extract word length from parafoveal preview information, allowing exact saccade targeting (e.g., Hawelka, Gagl, & Wimmer, 2010; Gagl, Hawelka, & Hutzler, 2014; Rayner, 2014; Gagl, Hawelka, & Wimmer, 2015; Meixner, Nixon, & Laubrock, 2022; for a more detailed discussion on the influence of visual noise, see Gagl et al., 2020). Furthermore, restricting the input to a specific word length also restricts the words in the lexicon that are used to generate the prediction (Gagl et al., 2020). Until now, the PEMoR assumed that all German words within the same word length are used to form the prediction. In a recent study investigating word recognition after learning, we knew the items of the lexicon (i.e., the words that were correctly perceived in most cases). We found that using the known lexicon items to calculate the prediction allowed modeling human and animal behavior in an orthographic decision task with high accuracy (Gagl et al., 2024). Thus, in this study, we explore whether we can increase the precision of the prediction error by restricting the number of words that are used to generate the prediction. We test if the model that includes only a subset of the lexicon results in a more adequate prediction error. Again, we use two variants, first restricting the words of a frequency-sorted lexicon (i.e., only the most frequent words are included) or a lexicon assumption that is not influenced by word frequency (i.e., randomly selecting the same number of words as frequency-based lexicon).

However, implementing a threshold or an adequate lexicon comes with the problem of selecting the most appropriate threshold value for the implementation and the appropriate set of words. We solve the issue by testing multiple PEMoR implementations, including varying thresholds and lexicon items for both new hypotheses (see example words in Fig. 1A, B). For model comparison, we test the variants against each other and also contrast the new implementations with the original *oPE* implementation. For model evaluation and comparisons, we use behavioral and electrophysiological brain data (see Gagl et al., 2020 for detailed descriptions).

## Method

### New Implementation of the prediction error model of reading

As assumed in PEMoR, the orthographic prediction error was implemented by image-based computations. For each pixel of the images (e.g., 5-letter words in Courier New font: 140×40 pixels in size), we estimate a knowledge-based prediction and stimulus-specific prediction errors (see Eq. (2)). Note that we simplify here. For the purpose of presenting the formulas, we recode the values 0 for black and 255 for white (standard in grayscale images) to black being represented by 0 and white by 1. Therefore, for both new hypotheses, the resulting prediction error per pixel can either have a high value of 1 when not predicted or a low value of 0 when correctly predicted. The difference between the two hypotheses is whether the binary algorithm was applied to the prediction, resulting in different model behaviors (see Fig. 1).

#### Mean prediction hypothesis

As shown in Figure 1A, the mean prediction is calculated in the same way as in the original implementation (Gagl et al., 2020), allowing a graded prediction error (i.e., parametric implementation with values between 0 and 1; see Eq. 1(1)).

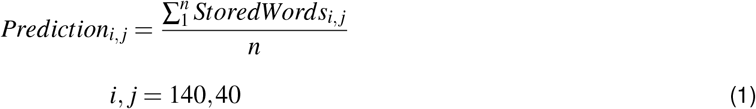

The prediction image (*Prediction*_*i,j*_) is the result of a pixel-by-pixel mean (i.e., *i, j* represents the pixel matrix of the images with 140 columns and 40 rows) over all images from all stored words (*StoredWords*_*i,j*_) in the lexicon (*n* represents the number of words in the lexicon). First, we implement a sum from all pixel matrices, conserving the position of the pixel in the matrix over all words of the lexicon. To calculate the mean for each pixel value, we divided the sum by *n* (i.e., the number of words in the lexicon). Note that the words used to approximate the lexicon (i.e., the stored words) for the prediction for each pixel were taken from the German SUBTLEX database (i.e., all five-letter words; Brysbaert et al., 2011). How we adapt the lexicon to achieve a more precise prediction is described below.

To achieve a more precise binary prediction error representation with a graded mean prediction, a threshold has to be implemented on the level of the prediction error (i.e., after the prediction is subtracted from the sensory input).

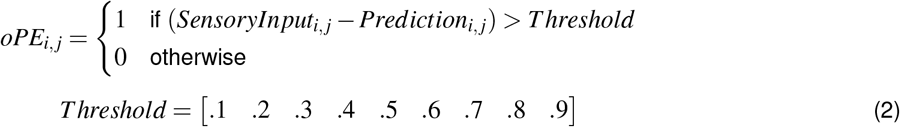

In this hypothesis, the resulting prediction error representation (*oPE*_*i,j*_) is achieved by first subtracting the mean prediction (*Prediction*_*i,j*_) from the sensory input (*SensoryInput*_*i,j*_) pixel by pixel. Note that the *SensoryInput*_*i,j*_ is a monochrome image with the value 0 when black and 1 when white. After that, we applied a threshold to all the pixel values, pixel by pixel (threshold values varied from 0.1 to 0.9, representing 10% to 90% thresholds). If the result of the subtraction was above the threshold value, the *oPE* pixel at that position was set to 1 (i.e., when the prediction was not correctly predicting the sensory input). If the result of the subtraction was below the threshold, we assumed a prediction error of 0 at the specific pixel(i.e., correctly predicting the sensory input). For example, with a .5 threshold (50%), when the sensory input of a given pixel is 1, and the prediction at that pixel is 0.2, the prediction error of that one pixel is 1 because the difference, 0.8 is higher than the .5. If the prediction is 0.6, the prediction error is 0 as the difference value, .2, is lower than the value of the threshold, .5.

#### Binary prediction hypothesis

Here, we assume a precise, binary prediction. Again, we first calculated the pixel-by-pixel mean of all the words in the lexicon. Different now is that we implement a threshold for each pixel value of the prediction matrix (*Prediction*_*i,j*_).

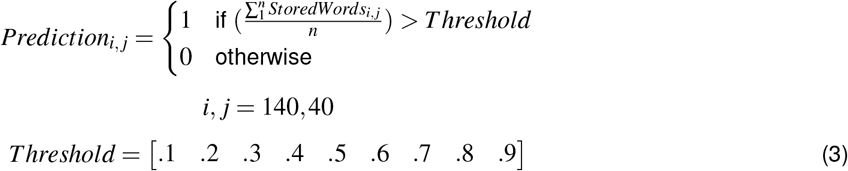

The pixel values of the binary *Prediction*_*i,j*_ (see Eq. (3)) are 1 when the average across the pixel values of the image representation of the *StoredWords* in the lexicon (i.e., *StoredWords*_*i,j*_) exceeded the *Threshold*. Thus, the pixel values of the binary *Prediction* can be a value that is either 0 or 1 (i.e., cp. the prediction images shown in Fig. 1B). A consequence of the binary prediction is that the prediction error that integrates a monochrome *SensoryInput*_*i,j*_ and a binary, therefore also monochrome, *Prediction*_*i,j*_ based on a subtraction can also only result in a monochrome *oPE*_*i,j*_ representation. Let’s look at the example from above. When the sensory input is 1, and the prediction is 0 (i.e., the mean prediction of 0.2 was set to 0 as the threshold at the prediction was higher with 0.5), the prediction error is 1 because the difference between 0 and 1 is 1. If the prediction is 1 (i.e., the mean prediction of 0.6 was set to 1 as the threshold was lower), the prediction error is 0 as the difference between 1 and 1 is 0. Thus, when implementing a threshold at the prediction or after the prediction error calculation (as in the mean prediction assumption), both result in a binary prediction error. Obviously, from the examples in Figure 1, the two assumptions result in very different *oPE*_*i, j*_ representations when thresholds are low or high. Still, already here, a higher similarity between the hypotheses at a threshold of 0.5 becomes visible.

#### Word frequency-driven prediction

To enhance the precision of the word lexicon, we rank all five-letter words from the German SUBTLEX database according to their frequency (i.e., the first word of the lexicon is the most frequent one and the last one the rarest word; see Coltheart, 2005; Brysbaert et al., 2011 for discussion on the implementation). We then generate predictions for chunks of words, ranging from the top 100 to the top 3108 highest-frequency words, in steps of 100. Accordingly, two prediction hypotheses were calculated separately for each of the 32 different word-frequency lexicons. In addition, as a control condition, we compared the frequency-ranked lexicons to a condition in which we randomly draw the same number of words multiple times (10 times for each same lexicon size as the frequency-based lexicon). These control conditions allow us to investigate if frequency sorting is essential for the lexicon.

### Model evaluations

For the model evaluations, we generated an *oPE* for each stimulus presented in the studies, separate for the two new and the original prediction error assumptions. In addition, one variant for each threshold and word lexicon assumption (*N* = 576 models). To create a parameter for model comparison, we summed up the pixel values from the *oPE* matrix for each stimulus. This summed *oPE* then represents the overall prediction error amount per word. So, as shown in the examples of Figure 1, the white values represent the amount of error per letter string (i.e., a large number of white pixels represents a high prediction error; see Gagl et al., 2020 for more details).

Here, we evaluate the new *oPE* implementations by comparing the model fit of regression models (i.e., linear mixed models). Specifically, we compare all 576 model implementations (i.e., hypotheses) with varying threshold levels and lexicon assumptions. For both hypotheses, the new *oPE* assumptions are compared separately for each lexicon. Additionally, we also compare these new *oPE* implementations to the original *oPE* model described by Gagl et al. (2020), assessing both prediction hypotheses under the same conditions. We use linear mixed effect models (Bates, Mä chler, Bolker, & Walker, 2014) and the Akaike Information Criterion (AIC, Akaike, 1973) to find the best fitting model. The behavioral data is based on a lexical decision task, and the brain data is based on electrophysiological measurements with EEG (see detailed descriptions of the dataset in Gagl et al., 2020). The EEG data gives us the chance to investigate neuronal implementation with a highly time-resolved dataset. Please note that the behavioral and EEG data are based on independent groups of participants.

#### Word Recognition Behavior

All stimuli of the lexical decision task (i.e., indicate by button press if the stimulus is a word or not), had five letters (800 Words & non-words), and participants were typically reading native speakers of German (*N* = 35; find the behavioral lexical decision data here: https://osf.io/d8yjc/). We estimated linear mixed models (Bates et al., 2014) for each of the *oPE* assumptions as predictors to describe logarithmic transformed response times (i.e., to account for the ex-Gaussion distribution of response times). Inspired by the previous analysis (see Gagl et al., 2020), we also estimated the interaction of the *oPE* and word lexicality (i.e., word, pseudoword or consonant strings), we also added decision accuracy, letter-string frequency within whole lexicon as covariates of no interest to the models. To account for variability across different letter strings, we included random intercepts for each unique letter string (see Eq. 4).

After initially fitting the linear mixed models across oPE assumptions generated from all word lexicons and different thresholds, to identify the optimal fit by accounting for variability across both participants and letter string comprehensively, we applied the ‘maximal’ model estimation methods (Bates, Kliegl, Vasishth, & Baayen, 2015) to assess the winning oPE models’ fit with all possible random effect components included. In this approach, we began by fitting the most complex model, which included all possible random slopes. However, due to the highly complex structure of random effects, these models often fail to converge. To address this, we incrementally reduced the model complexity by systematically removing random slopes. This stepwise reduction ultimately produced a convergent model formula that successfully fit the winning word lexicon and threshold oPE assumptions, as well as the original oPE generated from the corresponding word lexicon (see Eq. 5). For the model fit comparison, we used the Akaike Information Criterion (AIC; see Eq. 6) comparing the model including a *oPE* variant with the null model without an *oPE* variant (Akaike, 1973).

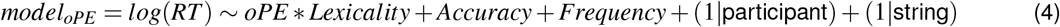

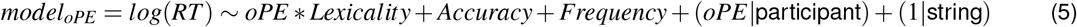

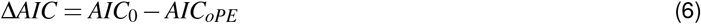

#### Electrophysiological data

To test the timing and effect changes of the new *oPE* on the neuronal level, we first focus on the two-time points previously identified as relevant (see (Gagl et al., 2020) for details): (i) posterior electrodes at 230 ms after stimulus onset showing an oPE effect and (ii) frontal electrodes at 430 ms after stimulus onset showing an interaction of the oPE with lexicality. EEG epochs were pre-processed (see Gagl et al., 2020), and for the first LMM-based analysis, in addition to the new oPEs, we include lexicality, string frequency and trial order as covariates of no interest. Random effects for participants and electrode locations account for variability due to individual differences and electrode characteristics (see Eq. 7). Again, the AIC compares the model fit (see Eq. 6). The ‘maximal’ model method was also conducted for oPE models on the EEG activation at two time points. When including random slopes, the model failed to converge for the original oPE at the 230 ms time window. Although the pattern between model comparisons is the same as using Eq. 7.

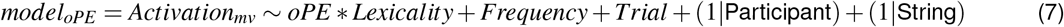

As an additional comparison, we again ran the linear regression analysis described in Gagl et al. (2020) for the entire EEG epochs (from 200 ms before and 800 ms after word onset). We used the multiple regression analysis with the same parameters as in the time-point-specific EEG evaluations and the winning oPE model (i.e., number of pixels, word/non-word, and the interactions with word/non-word distinction). Inference was made based on a cluster-based permutation test (Maris & Oostenveld, 2007). All clusters with a probability of less than an assumed alpha value of 0.05 under this simulated null hypothesis were considered statistically significant.

## Results

### Behavioral results

Model evaluation based on behavioral lexical decision data showed that the 50 % threshold models, both with mean and binary prediction, including the 500 and 1,800 most frequent words (i.e., 16% and 58% of the full lexicon; see Fig. 2A, B), had the best model fit. For both models, the fit was higher than the original oPE implementation when more than the 300 most frequent words were included in the lexicon. Applying the parsimonious mixed model approach to all four models, again, no difference was found between the algorithms. However, a higher model fit was identified for the 500-word lexicon implementation (see Fig. 2C). In addition, we ran 10 separate simulations in which we randomly drew either 500 or 1,800 words from the full lexicon to approximate an oPE version without a frequency-ordered lexicon (see Figure 2C). For all four models, we found that the frequency-ordered lexicon assumption for the oPEs estimation consistently provides a better model fit (500-word random lexicon: Mean AIC difference = 109.7; 95% confidence interval = [104.87, 114.53] vs. frequency ordered lexicon: AIC difference = 124; see Figure 2C).

**Figure 2:**
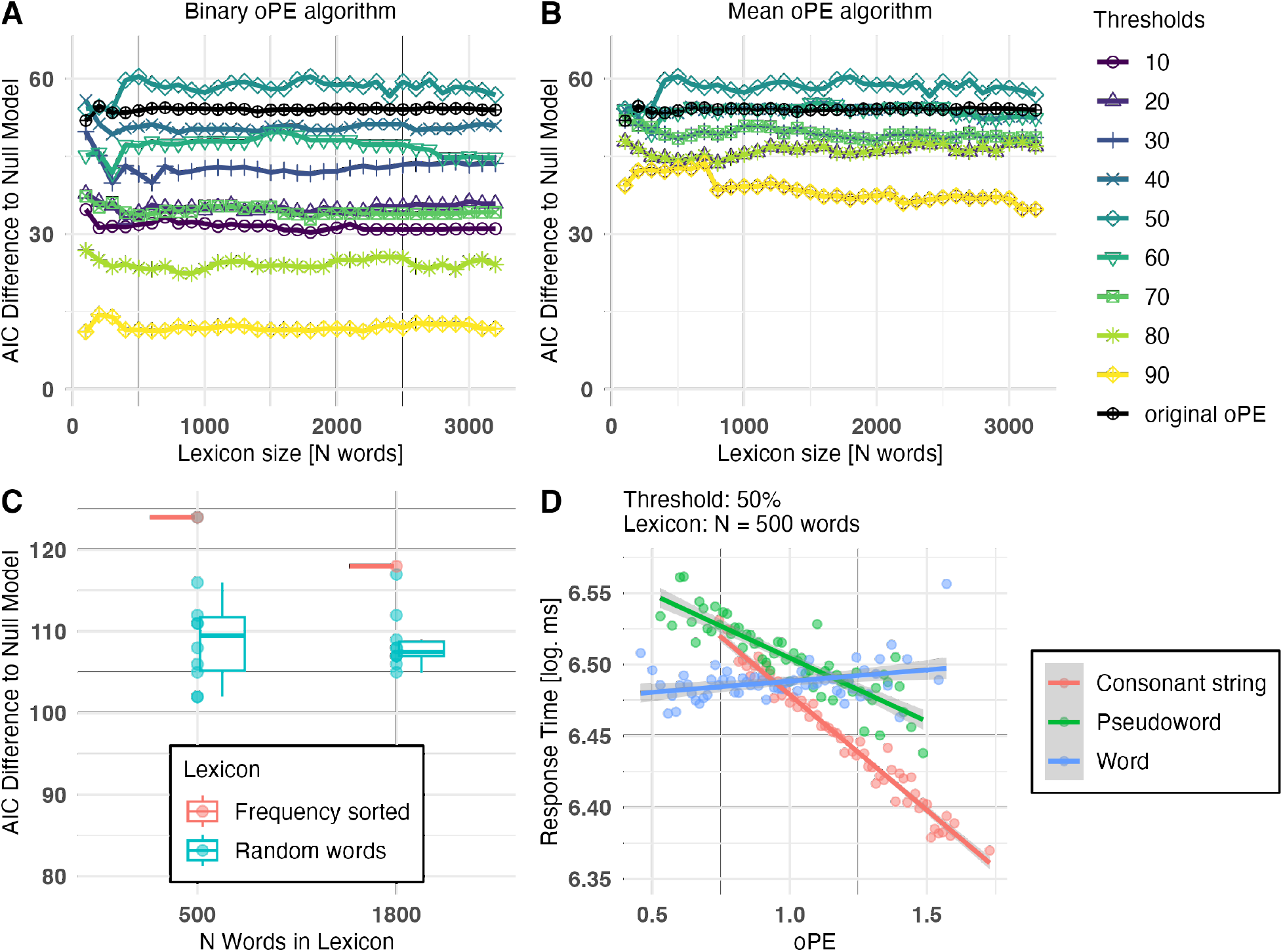
Model comparisons results for lexical decision behavior. (A) Model comparison results based on lexical decision response times for word frequency-driven oPE estimations, including all thresholds implemented on the level of prediction error under the binary prediction hypothesis. The x-axis represents the number of most frequent words used in the word lexicon to generate prediction and prediction errors each time. Model comparison results are also shown for original oPE (solid black line) generated from different word frequency word lexicons. All AIC differences presented here are compared against a version of the linear mixed model without an oPE predictor (Null hypothesis, H0). (B) Similar model comparison results are based on lexical decision response times for word frequency-driven oPE estimations, including all thresholds implemented on the level of prediction error under the mean prediction hypothesis. (C) Model comparison for randomly selected vs. frequency-ordered lexicons for the best fitting models (500 and 1800 word lexicons: 50% oPE). The results demonstrate that a frequency-based 50% oPE model using a 500-word lexicon provides a better model fit for human lexical decision performance. (D) 50% oPE from a 500-word lexicon affects response times in the lexical decision task. Blue lines show the effects for words, green lines for pseudowords, and red lines for consonant strings. Dots represent mean reaction time estimates across all participants, separated into oPE value and stimulus category after excluding confounding effects.

Inspecting the high similarity of the 50% threshold, the 500-word version of the binary and mean prediction models showed that they were perfectly correlated (*r* = 1; see Fig. 3A). For all other thresholds, we did not find such a high correlation between the two implementations. Furthermore, when inspecting the total prediction error, we see a linear decrease when assuming a mean prediction and an inverted U-shape function when investigating the binary hypothesis (see Fig. 3B). Therefore, we cannot differentiate between the two implementations, as the highest model converges to the only combination that does not allow a differentiation between the two hypotheses.

**Figure 3:**
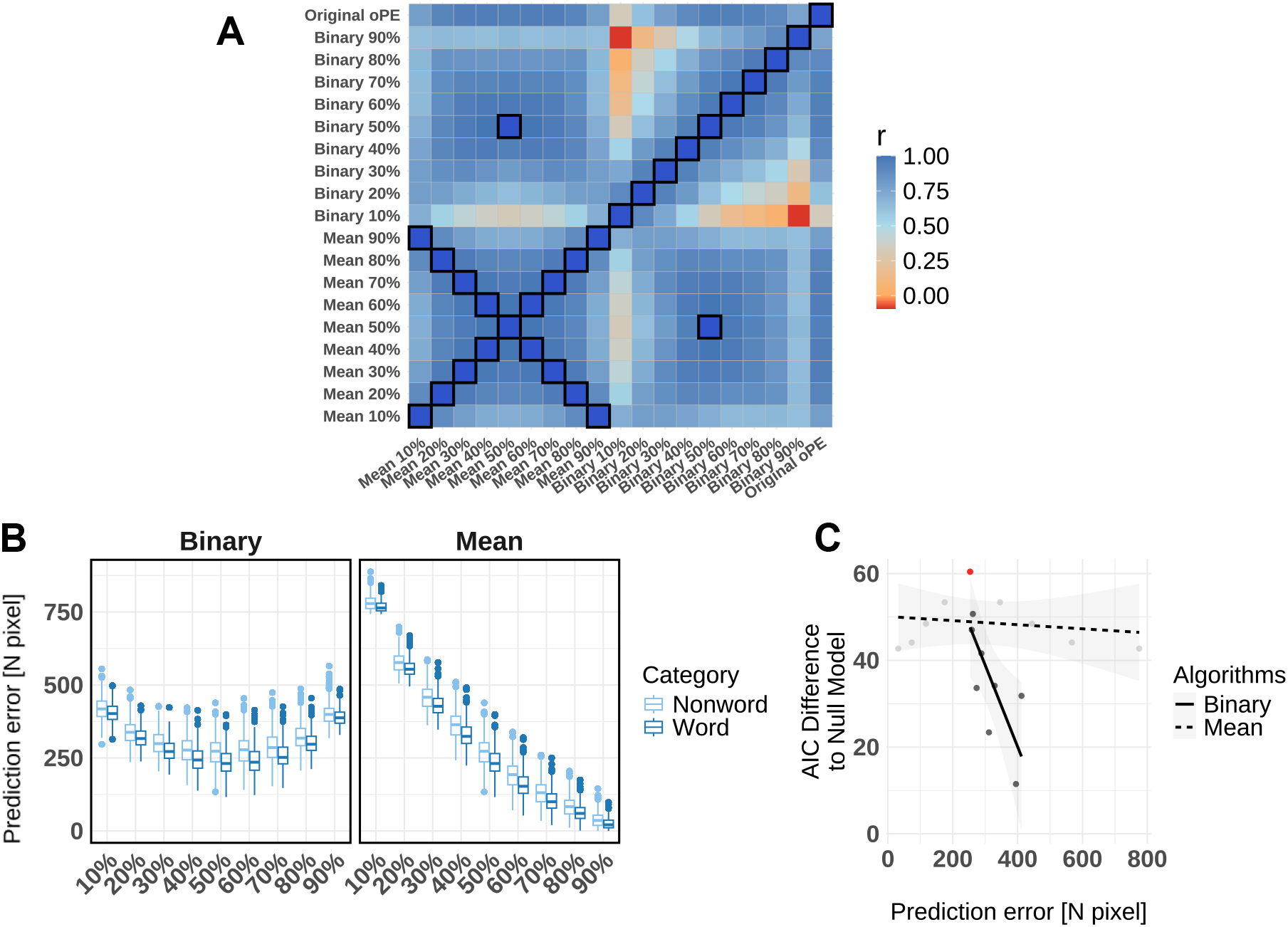
Similarities between and absolute oPE values of both algorithms and all thresholds when including the most frequent 500 words from the lexicon. (A) Correlation matrix between each oPE value from two algorithms at each threshold. All correlation coefficients range from -.09 to 1, with the perfect correlations marked by a black rectangle. (B) The number of error pixels of the oPE representation for each word or nonword (pseudoword and consonant strings) separated for all thresholds (x-axis) and both algorithms. Note that the number of error pixels is lower for words compared to nonwords. For the binary oPE, the difference at 10% is 16 pixels, which raises 42 pixels at the 50% threshold. The peak difference at 42.5 is found for the 60% threshold, which again declined to 11.5 pixels for the 90% threshold. For mean oPEs, the difference between thresholds is symmetric (i.e., similar to the correlations) with 15 pixels at 10% and 90%, 23 at 20% and 80%, 31 at 30% and 70%, 40 at 40% and 60%, and peaking at 42 pixels at 50%. Boxes and whiskers represent the distribution of the data, with the box spanning the interquartile range (IQR), which captures the middle 50% of the data, and the whiskers extend to 1.5 times the IQR, indicating variability outside the upper and lower quartiles. Outliers beyond this range are displayed as individual points. The black horizontal line represents the median of the distribution. (C) Correlation of the absolute oPE with the model fit separated by the algorithms. The red dot represents the best-fitting model (i.e., 500-word frequency sorted lexicon with a 50% threshold resulting from both algorithms).

Observing the 50% oPE on reaction time in lexical decision task based on the linear mixed model (see Fig. 2D), after excluding confounding effects, participants tend to quickly identify letter strings with low oPE value as words and the response is slower to recognize words with high oPE value; opposite pattern shown for pseudoword and consonant string categories, participants tend to respond slowly for low oPE value and quickly decide for letter strings with high oPE value.

### Simulation results

Inspecting the model simulations in detail (i.e., see Figure 3) gave a direct indication of why the model comparison could not differentiate between two implementations: (i) Mean and (ii) binary prediction algorithms with a 50% threshold. The two prediction error representations are perfectly correlated (see Figure 3A). Further inspection also revealed that absolute prediction errors resulting from the mean prediction algorithm showed high correlations between equidistant thresholds (i.e., 10 and 90% thresholds), which was not the case for the binary prediction algorithm. This finding can also explain why the model fits between equidistant thresholds from the mean prediction algorithm are equal (i.e., see overlapping lines in Fig. 2B).

Further, the absolute prediction error values showed a reduction of the prediction error values from low to high applied thresholds (see Figure 3B). When inspecting the prediction error example images in Figure 1A, one can see that with a low threshold, the error is large as the images include the errors that stem from the sensory input (i.e., the lower part of the “g”) but also errors that stem from the prediction (i.e., the errors in the area above the lower case letters). The prediction error decreases with increasing thresholds, resulting primarily from reducing the influence of the prediction on the error. In other words, from pixels where we expect that they are black in sensory input but, in contrast, are white instead. So, with a high threshold, only the errors that were not expected (i.e., not strongly included in the prediction) remained (i.e., the lower part of the “g”). The best fitting model, at the 50% threshold, results in a representation that holds both errors that stem from the prediction and errors that result from unexpected parts of the sensory input with an overall error that is in between the more extreme thresholds (see Figure 3B).

For the binary prediction algorithm, the 50% threshold results in the same errors, but here, the resulting errors are the lowest compared to all other thresholds. This finding indicates that the best-fitting model is also the model with the lowest errors in the representation, resulting in an inverted U-shape function (see Figure 3B). Also, we found that the correlation decreased between equidistant thresholds (see Figure 3A), indicating that the absolute error increases towards more extreme thresholds, but the represented information becomes more dissimilar. This finding stands in stark contrast to the implementation of the mean algorithm. When inspecting the examples of Figure 1B, one can identify the cause of this simulation pattern. The resulting prediction error representations are similar to the sensory input for high thresholds, a case not present in the mean algorithm. In contrast, with low thresholds, the errors only include the sensory input that is highly unexpected (i.e., the lower part of the “g”) plus the error that stems from the prediction—predicting away most of the sensory input, indicating that one can use the threshold in the binary prediction algorithm to navigate the importance of the sensory input or the prediction for the prediction error (e.g., high threshold - more sensory information; low threshold - more prediction information). Thus, that balance that we find at the 50% threshold represents both the information from the sensory input and the prediction in an optimal way with a low overall error.

Furthermore, when correlating the model fit indices (i.e., as shown in Figure 2A, B) with the absolute error (i.e., as shown in Figure 3B), we find a strong negative association for the binary algorithm (*r* = -.75; *t* (7) = −3.0; *p* = 0.02) but a weaker for the mean algorithm (*r* = −.19; *t* (7) = −0.5; *p* = 0.62; see Figure 3C). This finding indicates that the prediction error representation that results from the binary algorithm follows the pattern of the representation with the lowest error also holds the optimal representation. This relationship is not present in the mean algorithm. One further interesting observation is that the prediction error representation that primarily includes the sensory input (i.e., the 90% threshold from the binary algorithm; yellow line in Figure 2A) results in a lower model fit compared to prediction errors that are on the other more extreme end holding errors that largely stem from the prediction (i.e., low threshold condition of both algorithms). This finding replicates our previous finding that prediction error representation is a better representation when compared to the sensory input (i.e., see Gagl et al., 2020), but also that even prediction error variants with a high overall error, mainly holding the information included in the prediction result in a higher model fit (e.g., cp 90% and 10% in Figure 2A). Underlining the importance of prediction error representations for efficient visual word recognition behavior.

### Electrophysiological results

Based on the best model fit from the behavioral results, we adopted the new oPE formulations (50% threshold based on 500-word lexicon, mean, and binary implementations), which showed the best model fit response time. First, we analyze the two-time windows of central interest (230 and 430 ms after stimulus onset) identified when describing the original oPE implementation (Gagl et al., 2020). The original oPE had the highest model fit at amplitudes at 230 ms. The 50% oPE model (AIC = 990034) outperformed the null model (AIC = 990042) by 8 points. However, the original oPE model from the 500-word lexicon (AIC = 990033) performed even better, surpassing the null model by 9 points. Again, there is no difference between the mean and binary algorithm. At 430 ms, the 50% oPE model (AIC = 3185938) demonstrated an improvement over the null model (AIC = 3185940), with a 2-point difference. Additionally, the 50% oPE model outperformed the original oPE model from the 500-word lexicon (AIC = 3185939) by 1 point. Together, these results suggest that, at the early time window, around 230 ms, the original oPE was the best-fitting model. Still, at the later time window, around 430 ms, there is a consistent advantage of the 50% oPE model over both the null and original oPE models.

In addition, we investigated the effects of the 50% threshold meanbinary oPE based on the 500-word lexicon with a multiple-regression analysis for the full epochs. The models included the 50% threshold oPE (based on a 500-word lexicon), number of pixels, lexicality, and the interactions of the oPE with lexicality. In comparison to the original oPE, we again find a significant main effect of the new oPEs, but the cluster was only from 210 ms to 250 ms at similar sensors (cp. dashed lines vs. yellow area in Fig. 4A). This finding is in line with the 230 ms model comparison analysis, as we found a higher model fit and a larger cluster for the original implementation when compared to the new implementations.

**Figure 4:**
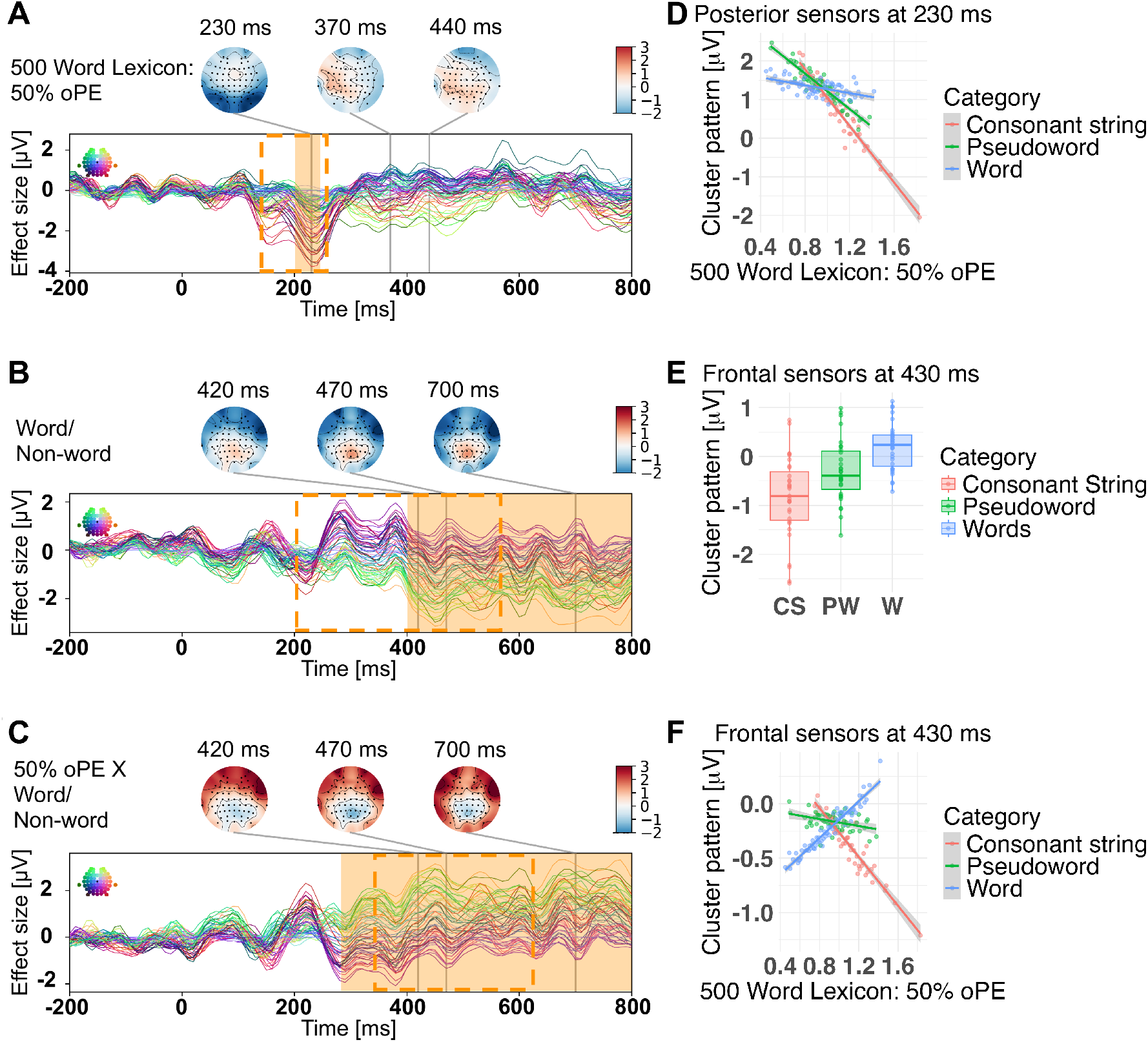
EEG results for lexical decision task of 200 words and 200 non-words (100 pseudowords, 100 consonant strings) including 50% oPE based on 500-word lexicon: Timing of 50% threshold based on 500-word lexicon orthographic prediction error effects separately. Effect sizes from multiple regression ERPs are presented as time courses for each sensor and time-point with orange shadow areas marking time windows with significant activation clusters (dashed rectangles mark the significant activation time windows from the multiple regression analysis for original oPE). ERP results are shown for (A) the main effect of the 50 % oPE based on a 500-word lexicon, (B) the word/non-word effect, and (C) the 50% oPE by word/non-word effects. The right panel shows the activation patterns related to the significant activation clusters in more detail. Dots represent mean predicted *µ*V across (D & F) all participants and items separated by 50% oPE and stimulus category, and (E) all items separated by stimulus category, excluding confounding effects. The frontal cluster includes the following sensors: AF3, AF4, AF7, AF8, F1, F2, F3, F4, F5, F6, F7, F8, SO1, SO2, FP1, FP2, Fz. The posterior cluster includes the following sensors: 02, 01, 0z, PO10, PO3, PO4, PO7, PO8, PO9, POz.

When inspecting the effect of lexicality, we found that with new oPE implementations, the cluster lasted from 410 ms to 800 ms. This finding, when contrasted to the lexicality effect in the analysis with the classical oPE implementation, started later in time but lasted longer (see 4B). The interaction cluster between 50% threshold oPE, and lexicality started earlier and lasted longer when compared to the original implementation. The cluster spanned from 300 ms to 800 ms and, therefore, even started before the lexicality effect cluster (see Fig. 4C). This finding changes the order of effects when compared to the original analysis from oPE cluster, lexicality cluster, and oPE by lexicality interaction cluster to oPE, oPE by lexicality interaction cluster, and lexicality cluster. This potentially indicates the change in the pattern of oPE effects from all negative to words positive and non-words negative, as shown in the interaction effect, before we access the words in the lexicon.

## Discussion

The Prediction error model of reading (PEMoR) highlights the role of orthographic prediction error representations (oPE) that integrate visual-orthographic knowledge with sensory inputs for efficient word perception (Gagl et al., 2020). Here, we revisited the original PEMoR and investigated if increasing the precision of the oPE is central to conserving critical visual information for efficient word identification. We explored adaptations of the original PEMoR formulation (two algorithms and 9 thresholds each), facilitating transparent theoretical development (i.e., as suggested by Guest & Martin, 2021), in which we increased the precision of the oPE on two levels: Graded vs. all-or-nothing (i.e., binary) oPE and full-size vs. partial frequency sorted lexicons. Overall, precision increases model fit. By implementing a binary oPE (i.e., precision by maximizing the differences between error and no error), pixel information becomes more easily differentiated (i.e., similar to a Quick Response code, QR code), increasing model fit. By implementing more realistic lexicon assumptions, the knowledge base of the model could be identified as a frequency-sorted lexicon containing a smaller number of words (i.e., as indicated by the word frequency effect; e.g., see Brysbaert et al., 2011; Gregorova, Turini, Gagl, & Võ, 2023), also increasing model fit. Overall, we found that both ways to increase precision resulted in adequate models describing visual word recognition behavior and late electrophysiological brain responses, outperforming the original PEMoR implementation. Only for early brain activation, we find the original oPE implementation to be the best-fitting model.

Our explorations identified the most optimal oPE implementing a threshold at 50% and a lexicon with only the most frequent 500 five-letter words (about 16% of all five-letter words in the SUBTLEX-DE, Brysbaert et al., 2011) resulting from both the mean and binary prediction algorithms. In the context of the current PEMoR implementation with five-letter words, 500 words would be roughly 16% of all five-letter words. In the context of an entire corpus (i.e., the 190,500 words in the SUBTLEX-DE), the 16% would translate to roughly 30,480 words, which aligns nicely with the estimated average adult productive vocabulary (i.e., about 30,000 to 40,000 words; e.g., see Brysbaert, Stevens, Mandera, & Keuleers, 2016).

Interestingly, both algorithms resulted in an oPE that perfectly correlated only for the implementation of a 50% threshold. This oPE best explains human behavioral performance and electrophysiological brain activation starting from 210 ms to 250 ms. In contrast, the original oPE implementation (i.e., graded prediction error) better described an electrophysiological component starting earlier from 150 ms to 250 ms. This pattern of results is intriguing as it indicates that one possibly implements precise prediction error signals in late processes (N400, behavior) but not necessarily in early processing. Using transparent computational models allows us to identify the differences explicitly and formulate a hypothesis on how the oPE representation changes from early to late time windows. The central difference between the original and the new formulation is that the individual pixel values are parametric in the former and binary in the latter, increasing the precision on the pixel level. The difference between early, graded prediction error and late binary prediction error could indicate a transformation of the oPE representation over time, settling on a solution with less nuance (i.e., only binary signaling). In other words, we might have identified a case of precision increase in a prediction error representation from early perceptual to later meaning retrieval processes.

Implementing the oPE representation as access code to word meaning (i.e., as proposed in Gagl et al., 2020), which becomes more precise over time, offers a simple algorithm that integrates sensory input and word knowledge for efficient lexical access. Similar to a QR code that stores, for example, the link to a webpage, our brain could implement the link between the visual stimulus of a word and its meaning via a binarised oPE representation. The central feature of the oPE representation, in contrast to a QR-code, is that it is not an arbitrary representation but based on a neuro-cognitively plausible assumption (i.e., predictive coding, Rao & Ballard, 1999) integrating the relevant information (i.e., word knowledge) with the sensory input while respecting the visual characteristics of the task (i.e., visual word recognition). Although optimized, it is still possible to infer the word when inspecting the oPE (see Figure 1). Therefore, when word knowledge is stored and predictive processing to optimize perception is implemented, one has all the ingredients to generate an oPE representation. Both are relatively widely accepted claims (see Coltheart, 2005; Norris & Kinoshita, 2012; Carreiras et al., 2014 for the assumption of word-knowledge; see Rao & Ballard, 1999; Alink et al., 2010; Price & Devlin, 2011; Clark, 2013; Heilbron et al., 2020 for the assumption of predictive coding). Still, a solution purely relying on visual information would be highly susceptible to various sources of error (i.e., through font changes or noisy sensory information), so it is highly likely that readers account for other sources of information (i.e., letters and letter-combination information) to, for example, make an access code more reliable (i.e., see Gagl et al., 2024 for a broader account of orthographic processing within a predictive processing framework). A technical solution for information storage, as implemented in the QR code, shares the implementation of a precise representation. However, it must implement arbitrary information irrelevant to the user (i.e., as the version of the encoding algorithm), being a reliable solution for the task. A barcode approach, as put forward by Hannagan, Agrawal, Cohen, and Dehaene (2021), based on a simulation study using deep-neuronal network computer vision models (i.e., a technical solution to visual perception, see van Rooij et al., 2023) is expected to have similar issues (i.e., one needs to code the model version or training regime). Thus, the oPE representation might be a pragmatic solution to the problem of representing sensory information in an optimal way within the parameters innate to the task of visual word recognition.

Interestingly, the binary and mean prediction assumptions for the oPE generation could not be empirically differentiated when focusing on the best-fitting model. The 50% threshold variant, irrespective of the number and type of words in the lexicon, was the best-fitting model but also resulted in identical oPE representations (i.e., perfect correlation). Nonetheless, the binary prediction and mean prediction algorithms could be differentiated when inspecting the simulations across all thresholds. Several aspects of both the binary and the mean prediction implementation are compelling, although the mean algorithm resulted in higher model fits over all thresholds. Also, the mean prediction algorithm can be best incorporated into the notion that brain activation evidence suggests a transformation from a parametric to a binary oPE representation. This is because both the original, parametric, oPE implementation and the binary oPE based on the mean prediction error algorithm implement the same prediction. Also, noteworthy aspects of the oPE representations from the mean algorithms are that irrespective of the threshold, strong prediction errors are always present, and the amount of the prediction error at a high threshold is lowest when compared to all other implementations.

The binary prediction algorithm, in contrast, is based on binary signaling only, making the algorithm more straightforward (i.e., binary signaling is less complex than the signaling of parametric information). Further, we find a correlation of the absolute prediction error (i.e., the sum of all error pixels) with the model fit, indicating that the best-fitting model also had the lowest prediction error compared to all other binary oPE implementations. The model fit decreases, and prediction error increases the lower the threshold gets. At low thresholds, the oPE representation becomes more similar to the sensory input (see Figure 1), including a high amount of redundant visual information. In contrast, when inspecting high threshold implementations, the prediction error is also increased. Also, we find that the high threshold oPEs primarily represent the information contained in the prediction (see Figure 1). The high dissimilarity of the oPE predictions resulting from low or high thresholds could be a meaningful feature one could test in future studies. As previously described in the implementation of the original oPE implementation (Gagl et al., 2020), visual noise in the sensory input changes the importance of the oPE for visual word recognition. At high noise levels, we even found that a representation of the sensory input resulted in a better fit for behavioral data. With this binary prediction algorithm, one could use the threshold as a parameter to adapt depending on environmental factors influencing the sensory input (i.e., sensory noise). Note that the latter is not inherent to the mean algorithm. Thus, a clear differentiation based on a direct empirical test is needed to specify the algorithm for the oPE implementation further.

The evaluation of new oPE formulations on EEG time courses shows that we can better explain early reading-related brain activation with the original oPE implementation than the new binary implementations. The oPE effect shares the time window and topography with the previously described occipitotemporal N170 and N250 components, which are both linked to early orthographic processing in visual word recognition (e.g., as they discriminate letters and words from non-linguistic stimuli; Hauk, Davis, Ford, Pulvermüller, & Marslen-Wilson, 2006; Barber & Kutas, 2007). Also, the components were associated with visual word characteristics (Dufau, Grainger, Midgley, & Holcomb, 2015; Sassenhagen, 2019), mark findings related to predictive processing from priming studies (Holcomb & Grainger, 2006; Grainger & Holcomb, 2009; Eisenhauer et al., 2019), and experiments with more natural sentence reading paradigms (Kornrumpf, Niefind, Sommer, & Dimigen, 2016; Milligan, Antúnez, Barber, & Schotter, 2023). Thus, the oPE effect, best described by an implementation of a graded error representation as described originally (Gagl et al., 2020), shows an effect in electrodes at a time window that was previously associated with prelexical visual-orthographic processes, typically preceding the lexical-semantic processes observed in later components.

From 300 ms post-stimulus, we find a significant interaction of the oPE with the lexical status of the letter string (i.e., word or nonword, lexicality) when using the best-fitting (on behavioral data) binary oPE (including a frequency sorted 500-word lexicon). About 100 ms later, we find a cluster showing the lexicality effect. Besides the earlier start of the interaction, both clusters overlap in time and topography. The change in the oPE from a parametric to a binary representation needs to be investigated in future work, but the pattern fits with the notion of a precise access code. The interaction effect could mark a process of accessing the lexicon before that lexicality effect marks access to the lexicon. Within this time range, the N400 component—typically associated with semantic and lexical processing—emerges (e.g., Lau, Phillips, & Poeppel, 2008; Kutas & Federmeier, 2011). Brain activation 400 ms after stimulus onset was associated with semantic congruity and expectation violations within reading, where it indexes the integration of word meaning and contextual fit within a sentence (Davenport & Coulson, 2011; Laszlo & Federmeier, 2011; Delogu, Brouwer, & Crocker, 2019) or lexical processing (Laszlo & Federmeier, 2009; Brothers, Swaab, & Traxler, 2015; Lau, Weber, Gramfort, Hämäläinen, & Kuperberg, 2016; Eisenhauer et al., 2019, 2022).Thus, the pattern of results aligns with theoretical frameworks in reading research that propose a transition from orthographic to semantic processing, with greater predictive precision enhancing the reader’s access to meaning (Hagoort, 2013; Kumari, 2022).

### Limitations

One of the central takeaways from the present study is that the oPE representation changes its form over time. This finding indicates a dynamic transformation algorithm to a currently static set of alternative models. This central limitation of the current approach motivates the implementation of a PEMoR version involving timing, so one can model the increase in precision before or while the oPE representation helps access word meaning. Further, the applicability of the oPE as an access code currently lacks font and size generalizability (i.e., at present, it is limited to a mono-spaced font that does not change in size) and focuses on the pixel level. In newer work, we expanded the idea of prediction error representations to the letter and letter-sequence level (Gagl, Weyers, & Mueller, 2021; Gagl et al., 2024), developing a generative model capable of simulating orthographic decision behavior across species. Still, future work must solve the issue of font and size generalizability, as humans can change the visual outlet of a text without detriment in reading efficiency. Changing the threshold parameter in the binary prediction algorithm could adapt the representation to different contexts, potentially allowing the adaptation to different visual configurations of a script. As noted before, this will be part of future research.

## Conclusion

Here, we revisited the PEMoR (Gagl et al., 2020), investigating if and how readers implement high precision in visual-orthographic representations underlying efficient word recognition. We found the implementation of a graded prediction error representation in early processing and a more precise prediction error representation in later processes (i.e., early vs. late brain activation) and behavioral responses. Especially for late processes, we found that higher precision in terms of visual reliability (i.e., graded vs. binary implementation of the representation) and lexicon structure (i.e., assuming that the lexicon consists of the 16% most frequent words) lead to more accurate model assumptions. Further, the new formulation provided insights into potential mechanisms to adapt the prediction error representation over time, as we found evidence of a change in the prediction error representation from a less precise graded to a more precise binary representation, potentially allowing more efficient access to meaning. Thus, our results provide a view on efficient visual-orthographic representations rooted in the visual information inherent in visual word recognition and the neuronally plausible predictive coding theory highlighting the role of error-based representations as key to accessing word meaning.

